# Long-Term Reconfiguration of Dynamic Polarization and Action Potential Initiation in Purkinje Cell Axons

**DOI:** 10.64898/2026.10.12.758508

**Authors:** Shin-ya Kawaguchi

**Affiliations:** Department of Biophysics, Graduate School of Science, Kyoto University, Oiwake-cho, Kitashirakawa, Sakyo-ku, Kyoto, 606-8502, Japan

## Abstract

Ramón y Cajal seminally predicted “the law of dynamic polarization of the neuron’’: synaptic inputs are integrated in the dendrites and soma, which are transformed into action potentials (APs) at the axon initial segment, faithfully propagating to the axon terminal for outputs toward target neurons. However, the exceedingly small size of axons and terminals in central neurons has hindered complete validation of this unidirectional signaling scheme. Here, using cutting-edge genetically-encoded fluorescent voltage imaging, complemented by tour-de-force axonal patch-clamp recordings, I show that the traditional one-way neuronal computation exhibits dynamic, activity-dependent long-term plasticity. In cerebellar Purkinje cells, AP firing, typically initiating near the soma, drastically shifts toward various sites of initiation in axons for more than two days, following the chemical induction of learning-related dendritic long-term postsynaptic plasticity. The dynamic shift of AP initiation is accompanied by enhanced GABA_A_ receptor activity and increased membrane excitability in the axon terminals. Thus, dendritic postsynaptic plasticity induces sustained axonal plasticity, giving rise to the emergence of local input-output elements at multiple terminals, which occasionally function as partially independent computation units. Together, the present study reveals an unprecedented dynamic modulation of neuronal processing, revolutionizing neuronal circuit computation.

## Introduction

Since Ramón y Cajal seminally proposed “the law of dynamic polarization,” neurons have been widely believed to integrate inputs in the somatodendritic compartment and then convey faithful digital outputs as action potentials (APs) along axons (Llinas, 2003; Abbott and Regehr, 2004). However, accumulating evidence is expanding this classical framework, revealing that axons function as digital-analog hybrid computational units (Debanne, 2004; Hoppa et al., 2014; Kawaguchi and Sakaba, 2015; Trigo, 2019; Trigo and Kawaguchi, 2023), where signals can be modulated by local neurotransmitters and APs can even be initiated locally at distal sites (Thome et al., 2018; Liu et al., 2022). A fundamental yet unresolved question is whether (and how) this multifaceted axonal computation undergoes activity-dependent long-term modulation or plasticity. Addressing this question has been historically hindered by severe technical hurdles: the minute dimensions of central axons and terminals (∼μm), their millisecond-scale kinetics, and their extensive trajectories (∼mm) that are typically severed in standard acute slice preparations.

To circumvent these obstacles, cerebellar Purkinje cells (PCs), the sole projection neurons of the cerebellar cortex, inherited with learning-related dendritic synaptic plasticity, serve as an ideal model system. I combined state-of-the-art genetically encoded fluorescent voltage imaging (St-Pierre et al., 2014; Hoppa et al., 2014) with subcellular direct patch-clamp recordings from boutons (Kawaguchi and Sakaba, 2015; 2017) in dissociated and organotypic slice cultures, where long axonal arbors remain intact, allowing for long-term functional evaluation. Here, I report that the chemical induction of learning-related dendritic postsynaptic plasticity causes a dramatic, sustained shift in the AP initiation site from the soma toward distal axonal terminals lasting for more than two days. This dynamic reconfiguration is enabled by enhanced axonal GABA_A_ receptor (GABA_A_R) activity and increased local membrane excitability, together with displacement of Na^+^ channels at axon initial segments (AIS). Together, my findings demonstrate that dendritic postsynaptic plasticity leads to long-term axonal plasticity, revealing a dynamic modulation of neuronal processing that fundamentally expands the computational capacity of individual neurons.

## Results

To detect neuronal activity at a high spatiotemporal resolution throughout a neuron, particularly aimed at detecting APs in axons, I improved a voltage-sensitive fluorescent probe protein, ASAP (St-Pierre et al., 2014; Vilette et al. 2019), by fusing the 72 (or even 30) amino acids of mGluR7 C-terminal (Stowell and Craig, 1999), a presynaptic membrane protein (Fig. 1A). Enhanced transportation of the probe towards the axon and terminals (Fig. 1B; here we denote it as axASAP) made it possible to locally detect AP arrivals at individual axon terminals, as well as synaptic inputs at any location of somatodendritic compartments, in PCs of dissociated and organotypic slice culture (Fig. 1; S1, S2, S3). Fortunately, axASAP unintentionally exhibited quicker fluorescence recovery after depolarization (Fig. S2), probably through changing physical status around the voltage sensor in the 4^th^ transmembrane segment, which enabled more reliable detection of even 100 Hz APs arrival at terminals (Fig. 1C). Interestingly, axonal voltage imaging highlighted not only APs but also slow transient depolarizing potentials at PC axons (Fig. 1D), resembling excitatory postsynaptic potentials (EPSPs), consistent with previous direct detection by axonal patch-clamp recordings (Zorrilla de San Martin et al., 2017). The axonal EPSPs (axEPSPs) were predominantly mediated by axonal GABA_A_Rs based on the higher axonal internal Cl^-^ concentration, as demonstrated by abolishment by picrotoxin (50 μM), a GABA_A_R blocker (see Fig. 2).

**Fig. 1.**
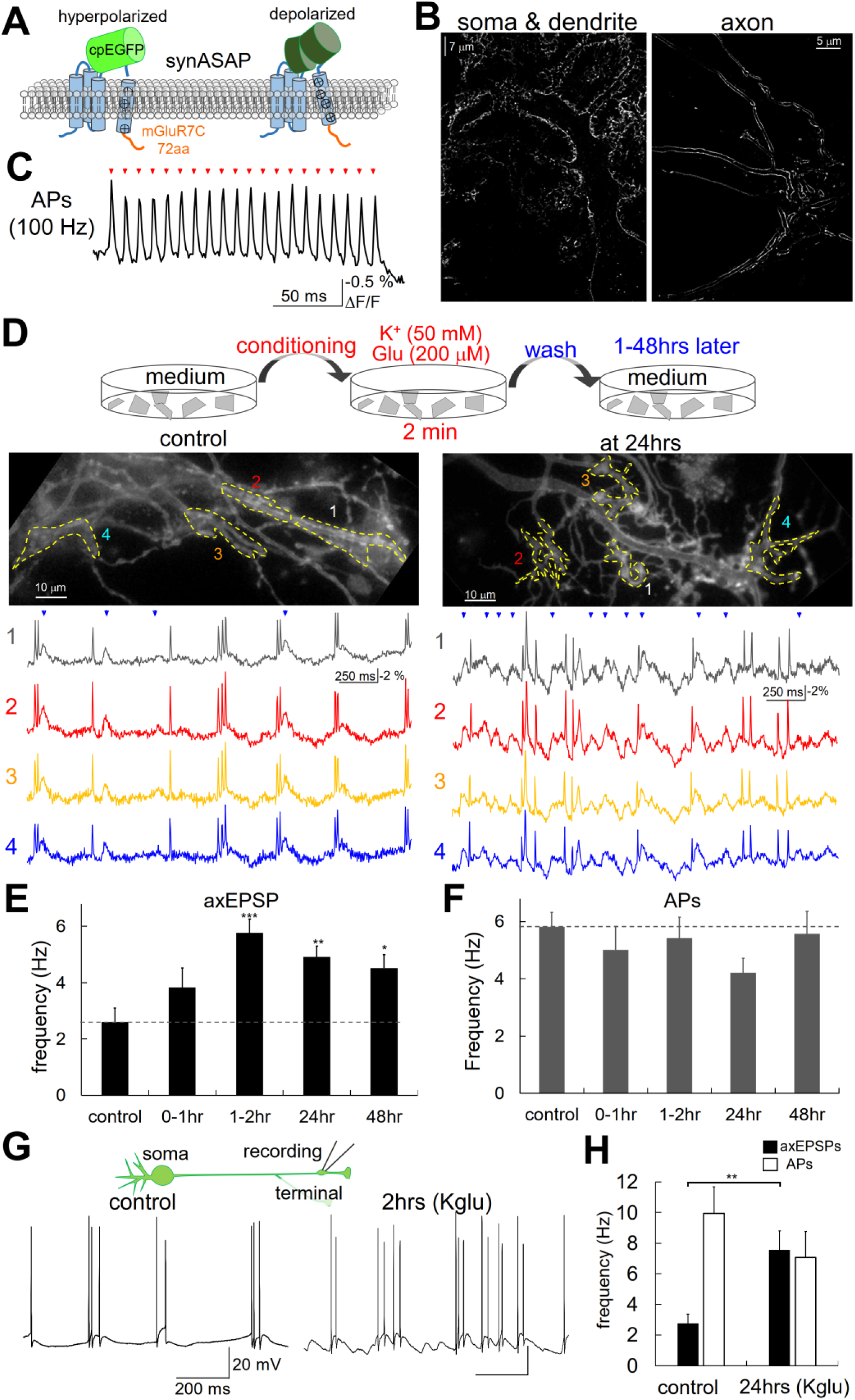
Voltage imaging of activity-dependent long-term increase of axEPSPs in a PC. **A**, **B** Schematic image (A), and representative SRRF confocal images (B) of improved ASAP (axonally-targeted ASAP: axASAP) at the soma/dendrite (left) and at a distal axon of a PC (right). **C**, Fluorescent imaging of 100Hz AP firing at axon terminals evoked at the soma. **D**, Top, Illustration of conditioning high K^+^ and Glu treatment for 2 min. Images of axASAP-expressing PC axon/terminals forming synapses on a target neuron before (middle left, control) and 24 hours after the K-glu treatment for 2 minutes (middle right, at 24hrs). ΔF/F traces at 4 different locations (bottom), showing spontaneous APs and axEPSPs. **E**, **F**, Frequency of axEPSPs (E) and spontaneous APs (F) before and after the Kglu treatment. N = 16 (control), 14 (0-1hr), 14 (1-2hr), 16 (24hrs), 14 cells (48 hrs). **G**, **H**, Representative traces (G) and quantification (H) of APs and axEPSPs directly recorded by a current clamp method from a PC bouton before and after the 2-min Kglu treatment. P < 0.01 in axEPSPs between control and Kglu. N = 8 boutons for both conditions. ***, p < 0.005; **, p < 0.01; *, p < 0.05.

**Fig. 2.**
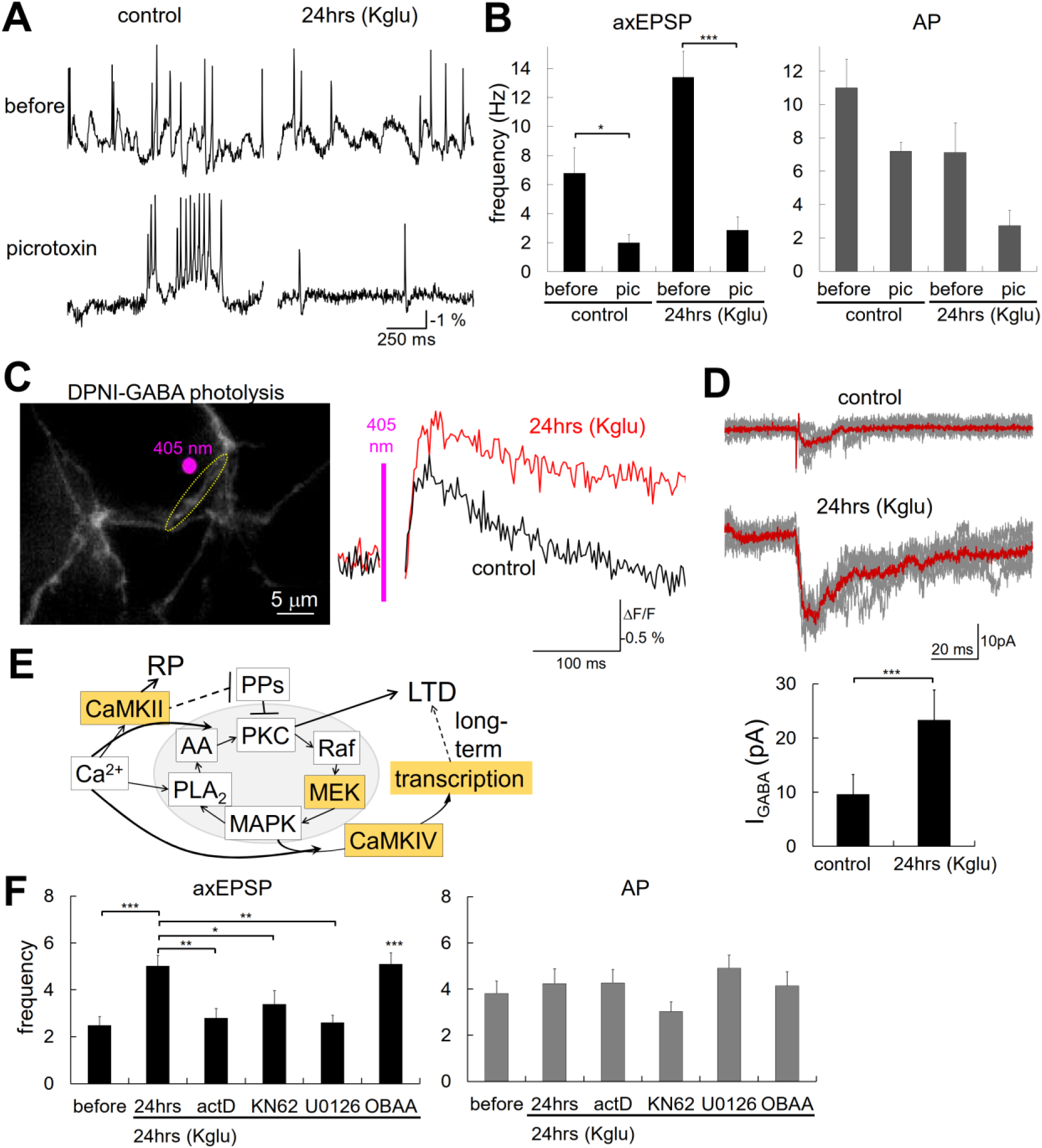
Enhanced axonal GABA_A_Rs underlying sustained plasticity of axEPSPs. **A**, Representative axonal Vm traces (A) and quantified frequency (B) of axEPSPs (left) and APs (right) recorded by voltage imagings before and after the picrotoxin application. N = 5 cells for each condition. **C**, Left, representative image of axASAP-expressing PC terminals stimulated by the 405 nm laser spot illumination (magenta) for uncaging of DPNI-GABA (300 μM). Right, fluorescently detected GABA-caused depolarization without (black) and 24 hours after (red) the 2-min Kglu treatment at PC boutons at roi (marked by yellow dotted circle in the left image). **D**, Direct axonal patch-clamp recorded traces (top) and amplitudes (bottom) of currents to locally photolyzed DPNI-GABA to an axonal bouton under before and 24 hours after the 2-min Kglu treatment. Red traces are the average of individual gray traces. N = 20 and 17 cells for control and 24hrs (Kglu), respectively. P < 0.01 by Wilcoxon test. **E**, Key signaling molecules involved in the induction of LTD/RP. Specific molecules tested are highlighted by orange. **F**, Frequencies of APs (right) and axEPSPs (left) observed by voltage imagings at PC axon terminals are shown in the control and 24 hours after the 2 min K-glu treatment with or without pharmacological agents. N = 17 (before), 21 (24hrs), 15 (actD), 17 (KN62), 15 (U0126), 18 cells (OBAA). ***, p < 0.005; **, p < 0.01; *, p < 0.05.

The axonal localization of ionotropic receptors which exhibit forms of postsynaptic plasticity in PCs (Kano et al., 1992; Kawaguchi and Hirano, 2000), made me assume that neuronal activity might also modulate those axonal receptors, which potentially impacts local computation in an axon. After treatment of neurons with high K^+^ (50 mM) and glutamate (200 μM)-containing external solution for 2 min (Murashima and Hirano, 1999; Kawaguchi and Hirano, 2013), fluorescent voltage imaging demonstrated that the axEPSPs, but not AP firing, became more evident in 1-2 hours, which persisted for longer than 2 days (Fig. 1D-F). Direct measurement of membrane potential by subcellular axonal patch-clamp recordings confirmed the long-term increase in events of axEPSPs at 2 or 24 hours after the brief K-glu treatment (Fig. 1G, H). Notably, the axEPSPs after the 2 min K-glu treatment were almost abolished by picrotoxin (Fig. 2A, B), indicating that the increased axEPSPs were mediated by axonal GABA_A_Rs. Fluorescent voltage imaging, coupled with local GABA stimulation by an uncaging of DPNI-GABA (300 μM) upon 405 nm laser spot illumination (∼2 μm in diameter) revealed augmentation of GABA-mediated local depolarization at PC boutons 24 hours after the K-glu 2-min treatment (Fig. 2C). Actually, the axonal currents to locally photolyzed GABA at the patch-clamped bouton nearly doubled after the brief K-glu treatment (Fig. 2D). Thus, the number and/or the function of GABA_A_Rs located at the axon/terminals are long-term augmented by the transient K-glu treatment.

Brief K-glu treatment typically causes intense Ca^2+^ increase in PC dendrites, leading to postsynaptic plasticity at excitatory and inhibitory synapses on PCs: long-term depression (LTD) (Ito and Kano, 1982; Ito et al., 2014) and long-term potentiation (LTP; specifically called as rebound potentiation, RP) (Kano et al., 1992; Kawaguchi and Hirano, 2000), respectively. Thus, I hypothesized that the long-term increase of axEPSPs is related to those forms of dendritic plasticity in terms of signaling cascades for induction (Fig. 2E). Pharmacological inhibition of Ca^2+^/CaM-dependent protein kinases by KN62 (1 μM), MAPK signaling by U0126 (10 μM) or *de novo* transcription by actinomycin D (20 μM), but not of PLA2 by OBAA (1 μM), during the 2-min K-glu treatment abolished the sustained increase of axEPSPs at 24 hours, but did not affect APs at terminals (Fig. 2F). These results indicate that the activation of plasticity-related transcription downstream of LTD/RP-inducing signaling cascades establishes the long-term increase of GABAergic axEPSPs (Tanaka et al., 2007; Tanaka and Augustine, 2008).

In addition to the augmented GABA_A_Rs, local membrane excitability reflected by the ratio of the voltage-gated Na^+^ conductance relative to the voltage-sensitive K^+^ conductance also became higher at PC axon terminals 24 hours after the brief K-glu treatment (Fig. 3A-C). Accordingly, immunocytochemistry showed increased localization of voltage-gated Na^+^ channels at terminals, but not at AIS, 24 hours after the 2-min K-glu treatment (Fig. 3D-F). Furthermore, interestingly, the Na^+^ channel distribution at the AIS near the soma got displaced away from the soma at 24 hours, keeping its length and total amount of Na^+^ channels (Fig. 3E-G), as reminiscent of the previous studies showing the activity-dependent re-localization of AIS (Kuba et al., 2006; 2010; Grubb and Burrone, 2010; Kole and Stuart, 2012). As a result of these various dynamic changes in axonal ionotropic GABA_A_Rs and membrane excitability, local stimulation of PC boutons by spot GABA photolysis became capable of triggering APs, back-propagations of which were detected at the soma by the cell-attached or whole-cell patch-clamp recordings (Fig. 4A-C). Activation of axon terminals by local GABA occasionally increased firings at the soma, on average into 2-folds for a few hundred milliseconds. As a result of such axonal plasticity, remarkably, voltage imagings of AP firings in axon terminals of an identical PC detected region-specific original AP firings which were not shared by the rest or some of other regions of the identical axon (Fig. 4D-F, S4, see red labels). Taken together, these results indicate that the augmented local GABA-sensing and the increased membrane excitability at PC axon terminals sometimes result in ectopic AP firings in an axon and/or terminals. Furthermore, such ectopic AP firing was occasionally specific to a restricted region of axon branches and/or terminals, leading to the emergence of potentially diverse output patterns among multiple terminals.

**Fig. 3.**
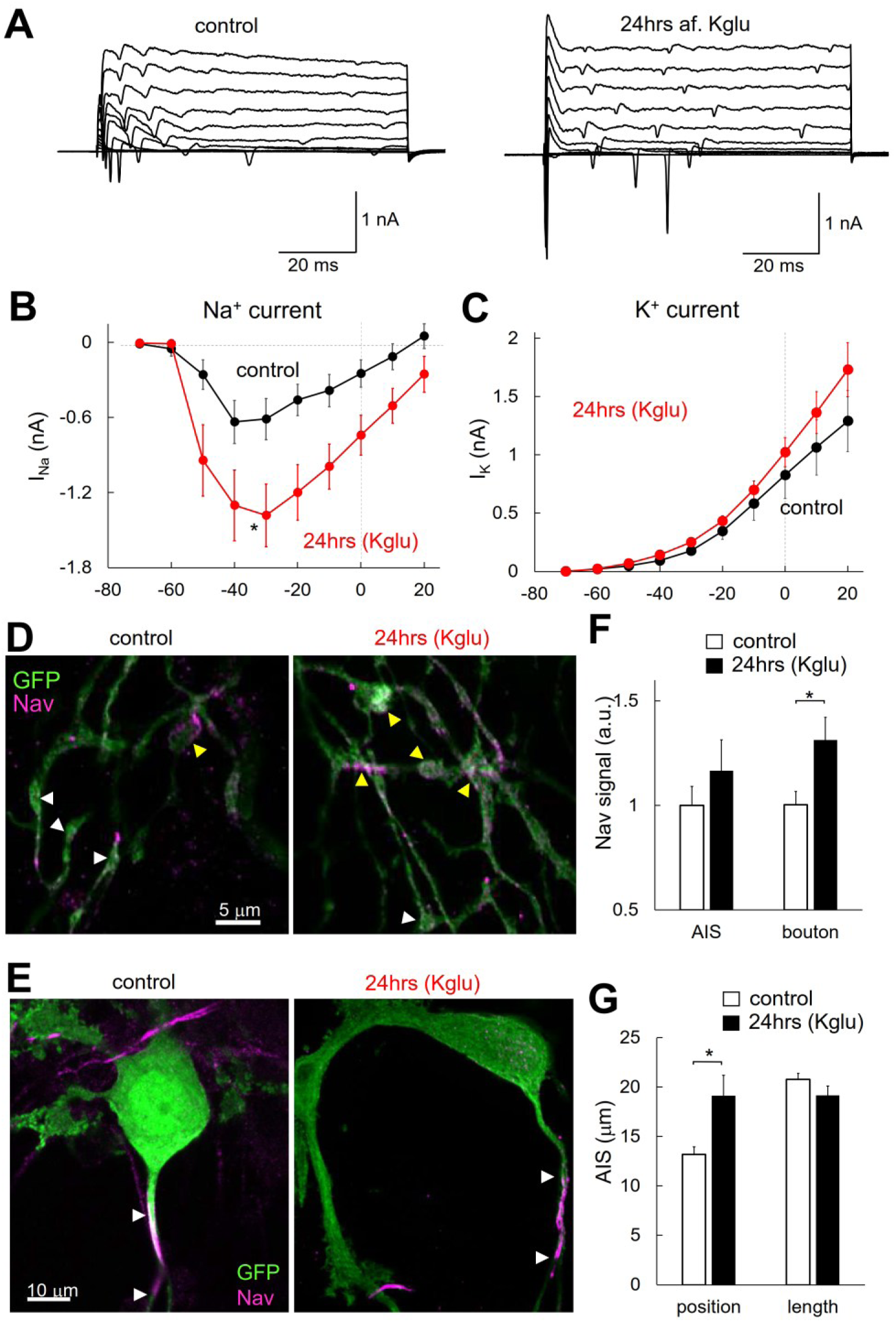
Enhanced bouton excitability associated with Kglu-triggered axonal plasticity. **A - C**, Representative traces (A) and I-V curves of voltage-dependent Na^+^ (B) and K^+^ currents (C) at a PC bouton recorded by direct patch-clamp recordings before and 24 hours after the K-glu treatment. N = 20 and 17 boutons for control and Kglu conditions, respectively. P < 0.05 by t-test (at – 30mV). **D-G**, Immuno-fluorescent images (D, E) of GFP (green) and Nav (magenta) in PC boutons (D) and axon initial segments (AIS) near soma (E) before and 24 hours after the K-glu treatment. Quantified Nav signal intensity and AIS length are shown in F and G. n = 40 and 39 boutons for Nav signal at AIS and bouton in control, and n = 41 and 39 boutons for Nav signal at AIS and bouton in Kglu condition, respectively. P < 0.018 at Nav signal between control and Kglu conditions. N= 40 and 41 boutons for AIS length in control and Kglu conditions, respectively.

**Fig. 4.**
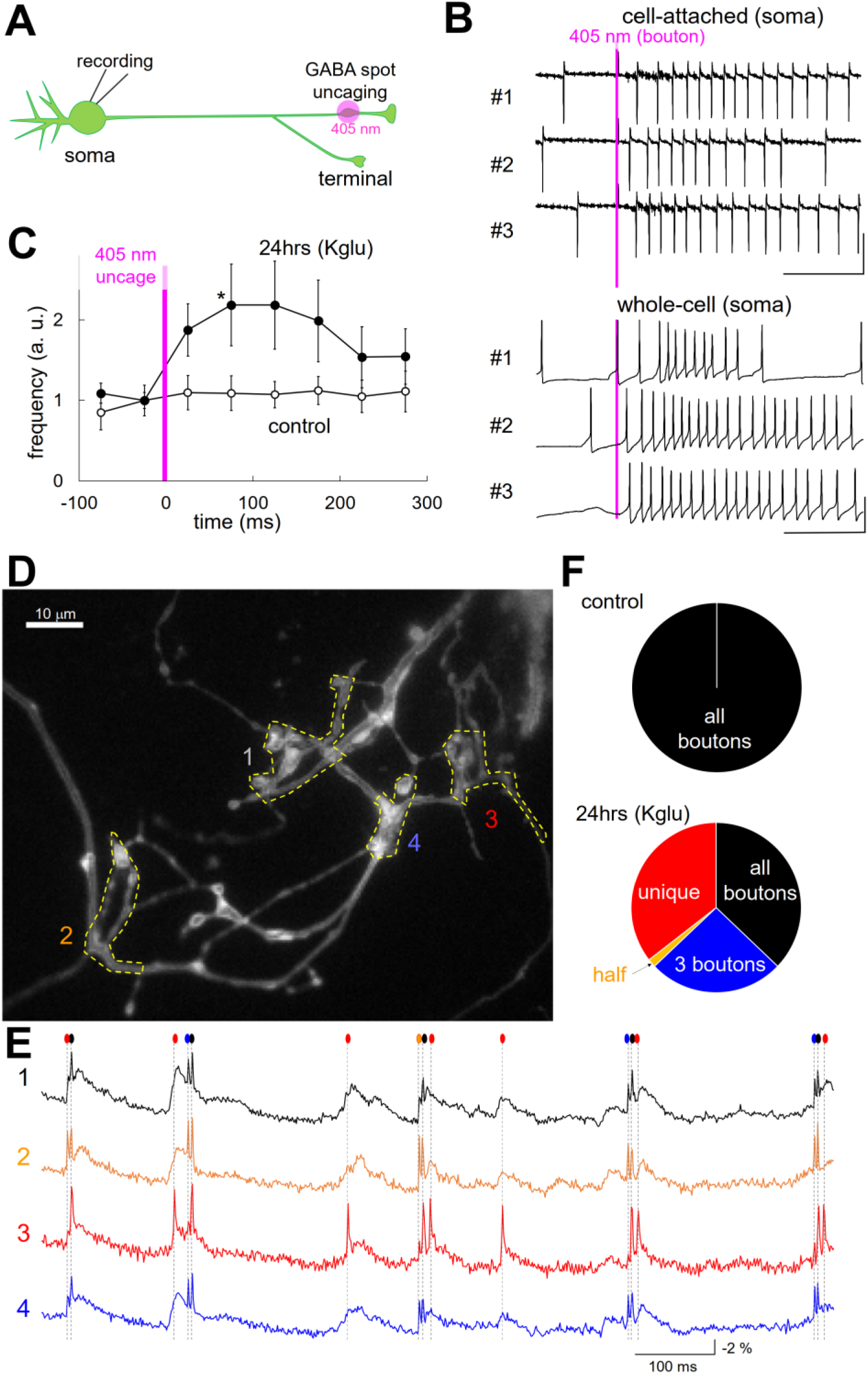
GABA-triggered firing initiation unique at PC boutons. **A**, Experimental design for somatic patch-clamp recordings of AP firings caused by local GABA uncaging at PC axon terminals distant from the recorded soma. **B**, AP recordings with somatic cell-attached (top) or whole-cell (bottom) configuration before and after GABA uncaging by 405 nm laser spot illumination at axon terminals in a PC 24 hours after the 2-min K-glu treatment. **C**, Frequency change of AP firings before and after the GABA uncaging in the control or K-glu treated PCs. **D**, Representative fluorescent image of ASAP-expressing PC axon/terminals of an identical PC 24 hours after the 2 min K-glu treatment. **E**, ASAP fluorescence traces at 4 ROIs (marked in D). Firings evident at all ROIs are marked by black symbol; only 3 ROIs, blue; only 2 ROIs, orange; unique at a single ROI, red. **F**, Pattern of common/unique firings at 4 areas in the analyzed field of view of PC axons/terminals before and 24 hours after the 2 min K-glu treatment.

To precisely evaluate how the AP firings initiate from distal axonal regions rather than the soma (i.e. AIS), I compared the timings of spontaneous APs at the soma and that at a distant (> 200 μm) axon terminal in a single PC, by performing dual patch-clamp recordings in cell-attached and/or whole-cell configurations (Fig. 5, S5, S6). In line with the Cajal’s law of dynamic polarization, APs basically appeared first at the soma prior to distal axon terminals (Fig. 5B, S5), in line with previous studies (Foust et al., 2010; Clark et al., 2015). The latency of AP detection between two sites showed almost single peak distribution (Fig. 5C, S5), which were comparable to the time needed for the somatically-evoked APs to reach the patched bouton (∼2 msec in average). The velocity of AP propagation along a PC axon from the soma to a bouton was ∼ 400-500 μm/msec, consistent with the earlier report for that from AIS to the axonal trunk in an acute slice (Clark et al., 2015) and in dissociated culture (Furukawa et al., 2024). These results altogether support the traditional view that APs predominantly initiate at AIS near the soma. However, intriguingly, APs recorded 24 hours after the plasticity-inducing brief K-glu treatment notably exhibited displacement and diversification of PCs’ firing sites: some of the APs recorded at the bouton preceded that in the soma, and *vice versa* (Fig. 5D, E, S6). The AP latency between the soma and the bouton exhibited a broader distribution, occasionally showing multiple peaks; the largest latency corresponded to the time needed for an AP triggered at the soma to reach the recorded bouton (Fig. 5F, S6). Intriguingly, the bifurcation pattern of length of an axon along the main path between the soma and the recorded bouton nicely overlapped the distribution pattern of AP latency. On average, the AP latency from the soma to a terminal significantly changed from positive to negative after the 2-min Kglu conditioning (control, 0.44 ± 0.24 ms; 24hrs (Kglu), −0.20 ± 0.16 ms, n = 12 and 16 PCs, respectively, p < 0.05, Student’s t-test). Thus, the plasticity-inducing K-Glu treatment caused dynamic change of spontaneous AP initiation sites in a PC axon, from the AIS near the soma to the multiple terminal regions beyond several axonal bifurcations. In addition, a subset of APs (∼ 17%) detected either in boutons or in the soma were absent in the other recording site after the K-glu treatment (8 out of 16 cells; Monsivais et al., 2005), supporting the idea that a fragment of axonal regions sometimes exhibits specific firing that does not spread to the rest of axonal branches (see Fig. 4). Inhibition of axEPSPs by GABA_A_R blocker picrotoxin tended to change the temporal distribution of AP latency in the K-glu treated PCs (Fig. S6), but not in control PCs (Fig. S5), although the tendency of ectopic firing initiation around axon terminal regions, not around AIS, was still remaining. Thus, the distal axonal firing property by itself seems ascribed mainly to the increased membrane excitability around terminals, and the local GABAergic inputs around terminals, if any, would trigger ectopic firings there. Finally, I examined whether the dynamic long-term change of AP initiation sites after the plasticity-inducing stimuli is the case for PCs *in situ*. While acute slice preparation was inappropriate for this evaluation because of the cut of axon during the slicing procedures, alternatively, organotypic slice culture preparation allowed me to study the AP initiation sites in an intact PC axon (Fig. 5G). As shown in Fig. 5H and 5I, qualitatively similar AP firings were observed in PCs in slice culture as those in dissociated culture: constant AP firings originating at the soma prior to the terminals in control, and the diverse and sometimes inversed order of AP occurrences at the bouton and the soma 24 hours after the 2-min K^+^-glu treatment.

**Fig. 5.**
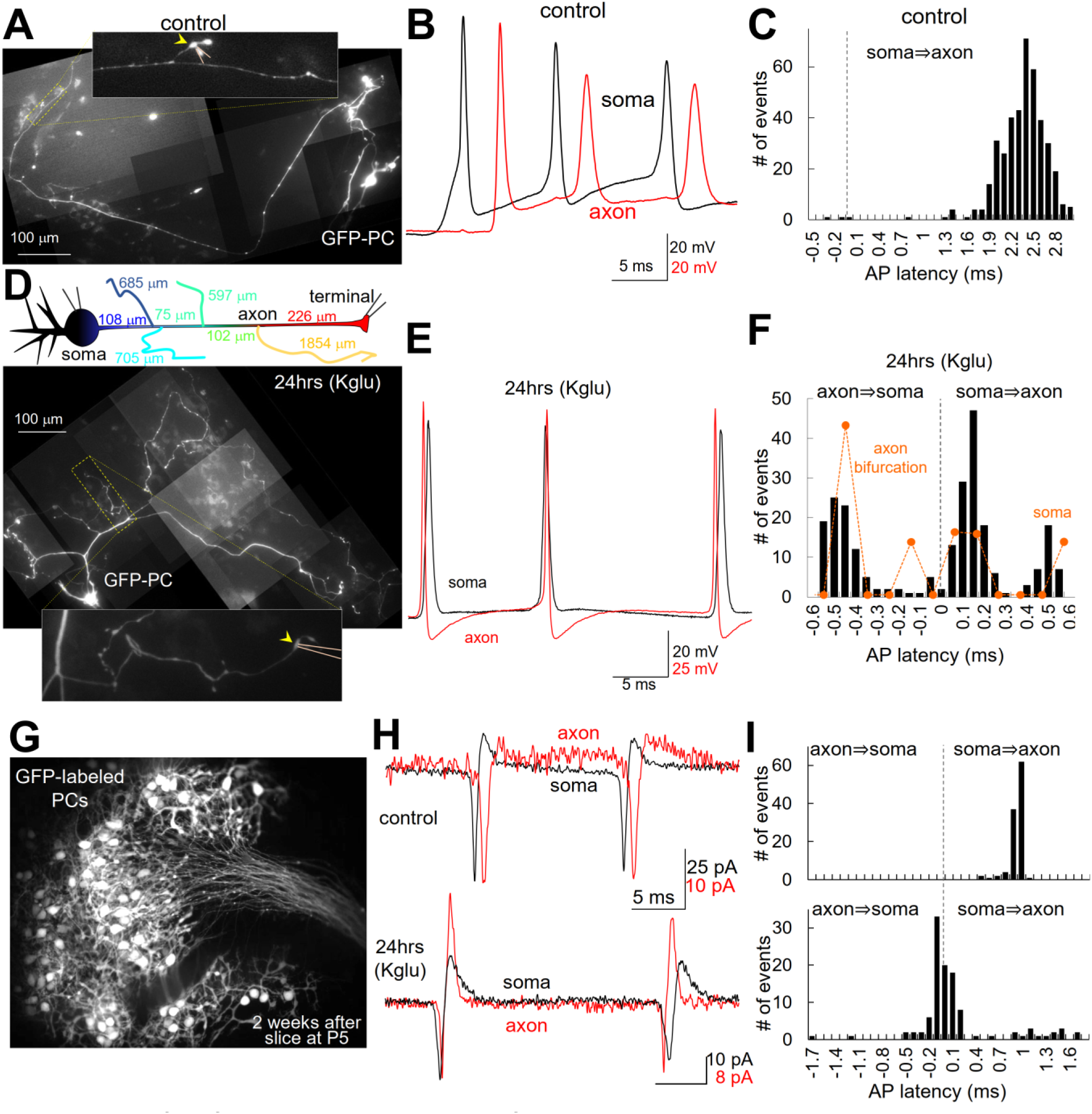
Diversified firing initiation in axons after the axonal plasticity. **A**-**F**, Fluorescent images (A, D), representative AP traces current-clamp recorded simultaneously from the soma (black) and the axon terminal (red, B, E) of a GFP-labeled PC in dissociated culture in the control condition (A-C) or 24 hours after the 2-min K-glu treatment (D-F). The histogram of AP latency between two sites (C, F). Schematic illustration of axon branching pattern is shown on the top of D, and the axonal area distribution is overlaid in F as an orange plot. **G**-**I**, Dual cell-attached AP recordings at the soma and an axon of a PC in slice culture in the control and 24 hours after the K-glu treatment. Representative GFP-labeled PCs (G), AP traces (H) and its latency between two recording sites (I) are shown.

## Discussion

By leveraging cutting-edge fluorescent voltage imaging and subcellular patch-clamp recordings from axons and terminals of PCs, I performed high spatiotemporal analysis of AP initiations and subthreshold membrane potential changes, revealing that the neuronal computation fundamentally diverges from the historical dynamic polarization theory presented by Cajal: 1, the distal axon and terminals are receptive for GABAergic inputs from autaptic and/or heterosynaptic neuronal sources (Trigo et al., 2010; Zorrilla de San Martin et al., 2017), which sometimes leads to local trigger of firing there, as structurally conceptualized in cortical axo-axonic microcircuits (Szabadics et al., 2006); 2, neuronal activity such as one inducing synaptic plasticity at dendrites results in long-term change of the AP initiation site from the somatic area to the distal axonal regions through dynamic rearrangement of axonal excitability and GABAergic sensitivity. Although ectopic AP firing initiation at axons was reported in hippocampal networks (Colbert and Pan, 2002; Dugladze et al., 2012; Epsztein et al., 2010), it has generally been regarded as abnormal activity associated with excitotoxicity or epilepsy, and its drastic impact on computation in neuronal network has not been conceptualized as a physiological computational mechanism. As demonstrated in the present study, ectopic axonal firing is subject to fine-tuning through activity-dependent tight control of axonal membrane excitability and local ionotropic receptors. While APs faithfully propagate across axonal bifurcations under basal conditions (Foust et al., 2010), our findings demonstrate that bifurcations with long axonal branch can be transformed into initiation nodes in the main axonal path. Thus, the dynamic modulation of neuronal compartments sometimes enables AP firing in a restricted subpopulation of an axon, giving rise to emergence of small input-output elements inside a single neuron.

Considering the cerebellar network, many studies have shown that motor learning is brought about by the activity-dependent modulation of information processing in the cerebellar cortex, through combinations of several forms of synaptic plasticity including LTD at excitatory synapses on a PC (Ito et al., 2014). However, curiously once established, the motor learning memory no longer relies on the activity of cerebellar cortical neurons, implying the presence of a mysterious mechanism for the transfer of memory trace encoding the motor learning from the cortical synapses towards the deep cerebellar nuclei (DCN; Wada et al., 2007; Okamoto et al., 2011). The long-term change of PCs’ firing sites from the somatic compartments towards the axon terminal regions might contribute to such a mysterious long-distance memory transfer, by producing local input-output elements in the DCN regions after the learning-related plasticity induction at PC dendrites. Taken together, the dynamic long-term plasticity of axonal information flow unveiled here fundamentally enriches our understanding of how neuronal networks communicate and process information.

## Competing interests

Authors declare no competing interests.

## Data and materials availability

All data are available in the main text or the supplementary materials.

## Acknowledgement

I am grateful to Kawaguchi Lab members for critical reading and constructive discussion for this project and manuscript.

## Funding

This work was supported by the grants from Japan Society for Promotion of Science, KAKENHI grants 22H02721, 22K19360, 25H02611, 25K02362, 26K21905 and Takeda Science Foundation.

## Experimental Procedures

### Animal usage

For preparing dissociated and organotypic slice culture, Wistar rats (P0 for dissociated and P5-6 for slice culture, respectively) were decapitated in accordance with the guidelines regarding care and use of animals for experimental procedures of the National Institutes of Health, U.S.A. and Kyoto University, and approved by the local committee for handling experimental animals in Kyoto University.

### Dissociated and organotypic slice culture

The method for preparing primary dissociated cultures of cerebellar neurons was similar to that in a previous study (Kawaguchi and Sakaba, 2015). For slice culture, rat sagittal brain slices (250-300 μm thickness) of the cerebellum were prepared from P5-6 pups with a Mcllwain tissue chopper (MK-MTC). The slices were kept on Millicell membrane (Millipore) mounted in the BME-based culture medium identical to that used for dissociated culture. PCs were transfected with EGFP or voltage-sensitive fluorescent protein modified from axASAP at 1 day after culture with an adeno-associated virus (AAV) vector under the control of CMV promotor (Kawaguchi and Sakaba, 2015). PCs could be visually identified by their large cell bodies and thick dendrites, and fluorescent protein which preferentially labeled PCs by relatively specific AAV serotype-2 infection. An axon of PC was clearly different from dendrites which have high density of spines. Whole-cell patch clamp recordings, fluorescent imagings, and immunocytochemistry were performed 3-4 weeks after preparation of the culture.

### Construction and transfection of DNA vectors and AAVs

The coding DNA sequence for ASAP1 was obtained from Addgene (#52519), and some modifications for higher voltage sensitivity (D145D, L146G, S147T, N149R, S150G, and H151D) established in an improved version ASAP3 (Villette et al., 2019) were introduced by PCR-mediated mutations (noted here as ASAP3β; Higashi et al., 2024). In addition, to enhance the intracellular trafficking of the ASAP protein toward axons and terminals, 30- or 72 mer C-terminal region of mGluR7b, which was obtained by RT-PCR of cDNA template prepared from rat cerebellum, was fused to the C-terminal of ASAP at the Bgl II site. The ASAP cDNAs were inserted into pCAGplay DNA expression vector (Kawaguchi and Hirano, 2013) and/or pAAV-CMV-WPRE vector prepared from pAAV-CMV (Takara) with addition of WPRE sequence immediately after the poly-A addition signal region. AAVs for types of ASAP were prepared by co-transfection of HEK 293T cells with an axASAP-encoding pAAV-CMV-WPRE plasmid, pHelper plasmid, and pAAV-RC2 plasmid, followed by collection of AAV particles 3-4 days after the co-transfection using AAV purification kit (Takara). Typically, the AAVs had titer of 10^9^ –10^10^ (particles/ml) and preferential infection of AAV2 to PCs among neurons of cerebellum, enabled specific transfection of ASAPs in PCs.

### Membrane potential fluorescent imaging

Original or modified ASAP probes expressed in PCs or HEK cells were illuminated (15 mW/mm^2^) by 450-480 nm blue light (Light Engine SOLA, Lumencore), and the fluorescent images were obtained by a sCMOS camera (Zyla4.2, Andor) with a x60 optical lens (NA 1.42) mounted on Olympus IX71 at 400 (512 x 512 pixels) – 1500 (128 x 128 pixels) frames per second. Cells were kept in the normal external solution containing (in mM): 145 NaCl, 5 KOH, 2 CaCl_2_, 1 MgCl_2_, 10 Hepes, and 10 glucose (pH 7.3) at room temperature (20∼24 °C). Fluorescent images were analyzed with ImageJ and the time course of fluorescent intensity was analyzed with Microsoft Excel and Igor Pro8 (WaveMatrics). In some experiments, GABA was locally applied by the photo-activation of DPNI-GABA (300-500 μM, Tocris Cookson, UK) with spot illumination of 405 nm laser (2 mW for 0.5 msec, OBIS, Coherent) at the axon terminal region of PC as in a previous study (de San Martin et al., 2017). Fluorescent imagings were performed from PCs without or after the 2-minutes conditioning treatment with an external solution containing 50 mM KCl (in place of NaCl) and glutamate (200 μM), followed by wash twice. In some experiments, PCs were pre-treated with KN62 (1 μM), U0126 (10 μM), OBAA (1 μM), or actinomycin D (20 μM) for 1 hour before the conditioning.

### Electrophysiology

Methods used for electrophysiological experiments were similar to those in previous studies (Kawaguchi and Sakaba, 2015). Whole-cell patch clamp recording from a cultured PC was performed with an amplifier (EPC10, HEKA, Germany) in the normal external solution at room temperature (20∼24 °C). In some experiments, 2,3-Dioxo-6-nitro-1,2,3,4-tetrahydrobenzo[f]quinoxaline-7-sulfonamide (NBQX, 10 μM, Tocris, UK), tetrodotoxin (TTX, 1 μM, Wako, Japan), picrotoxin (50 μM, Sigma) were used to inhibit glutamatergic EPSCs, action potentials, respectively. A patch pipette used to record from the soma, axon, and the terminal of a PC was filled with an internal solution (pH 7.3, adjusted by KOH) containing (in mM) 155 KCl, 0.5 EGTA, 10 Hepes, 2 Mg-ATP and 0.2 Na-GTP. The membrane potential of a PC was held at –70 mV. Series resistances of the PC soma and axon terminal were ∼10 and 70 MΩ, respectively. Series resistance of the terminal was corrected by 30-60 %.

### Immunocytochemistry

Cultured neurons were fixed with 4 % paraformaldehyde in phosphate buffered saline, permeabilized with 0.5 % Tween 20, then blocked with 2 % horse serum, and finally labeled with primary and secondary antibodies. The following antibodies were used: a mouse monoclonal antibody against Na^+^ channels (1 : 1000, Sigma), a chick polyclonal antibody (pAb) against GFP (1:1000, Chemicon), and CF 488A or 633-conjugated pAbs against rabbit or chick IgG (1:400, Biotium). The fluorescence images were obtained with the Dragonfly rapid confocal imaging system (Andor) mounted on an inverted microscope IX83 (Olympus), and analyzed with Image J software (NIH).

### Statistics

Data are presented as mean ± s.e.m. Statistical significance was assessed by paired t-test, unpaired Students’ t-test, Mann-Whitney U test, or ANOVA and the following Dunnett’s test.

---

**Supplementary Fig. 1.**
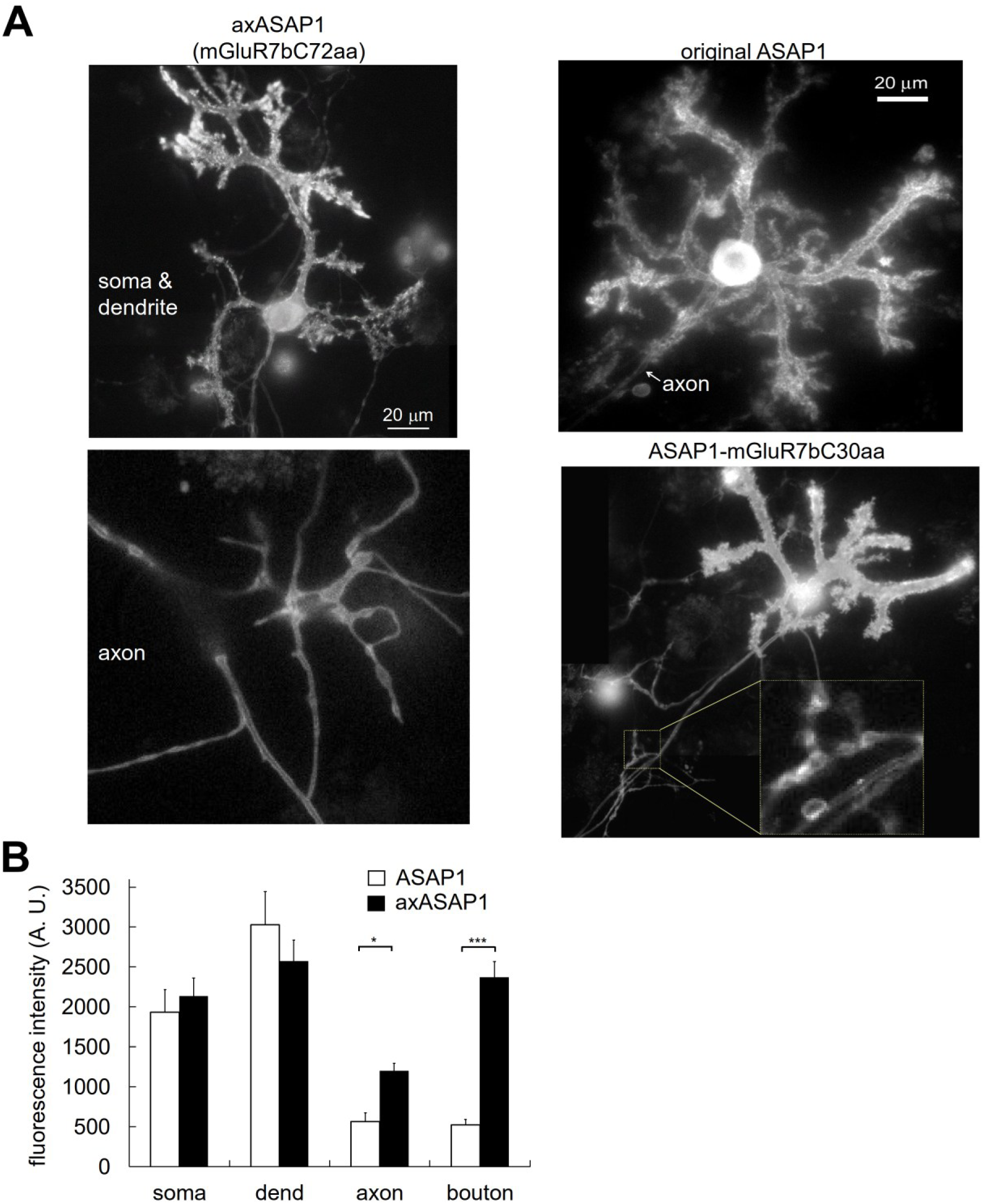
Subcellular distribution of ASAP. **A**, Representative SRRF confocal images of improved ASAP (axonally-targeted ASAP: axASAP) at the soma/dendrite (left) and at a distal axon of a PC (right). **B**, Fluorescent intensity at soma, dendrite, axon, and presynaptic bouton of original ASAP1 or axASAP. N = 41 (ASAP1) and 39 cells (axASAP1). ***, p < 0.005; *, p < 0.05.

**Supplementary Fig. 2.**
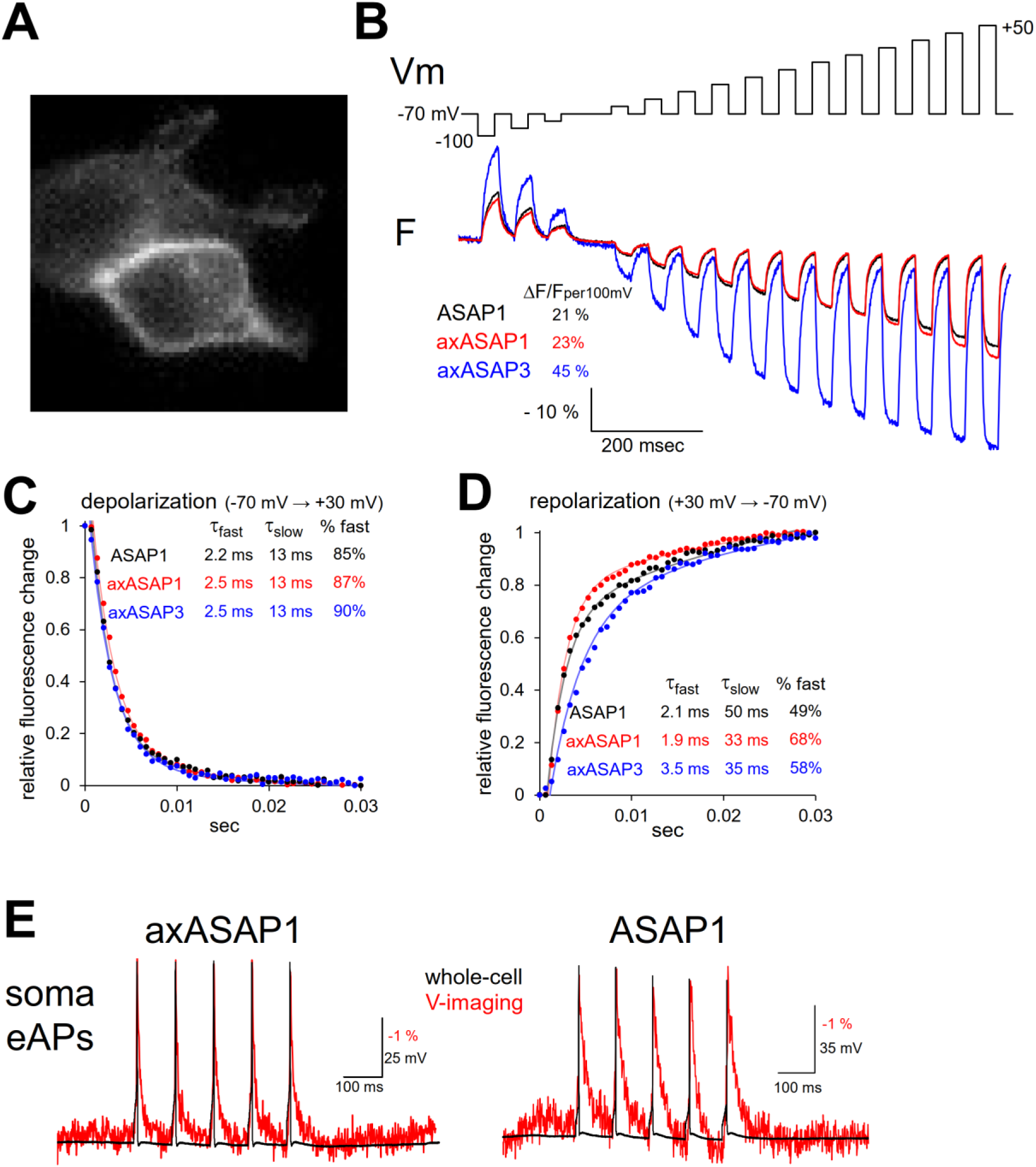
Biophysical characterization and fast optical kinetics of axASAP. **A**, Representative image of a HEK293T cell expressing axASAP3. **B**, Voltage-step protocol (holding potential: −70 mV; steps to −100 ∼ +50 mV) and corresponding optical fluorescence responses (ΔF/F) of ASAP1, axASAP1, and axASAP3. **C**, **D**, Normalized fluorescence change time course during 100-mV depolarization (C) and repolarization (D), comparing original ASAP1, axASAP1, and axASAP3. Fast (Ƭ_fast_) and slow (Ƭ_slow_) time constants and the relative amplitude of the fast component (% fast) are indicated for each probe. **E**, Simultaneous fluorescent voltage imaging (red, 1.5 kHz) and whole-cell current-clamp electrical (black) recordings of a single AP in the PC soma expressing axASAP1 (left) or original ASAP1 (right), illustrating the fast fluorescence recovery of axASAP1 providing better fluorescent tracking of APs.

**Supplementary Fig. 3.**
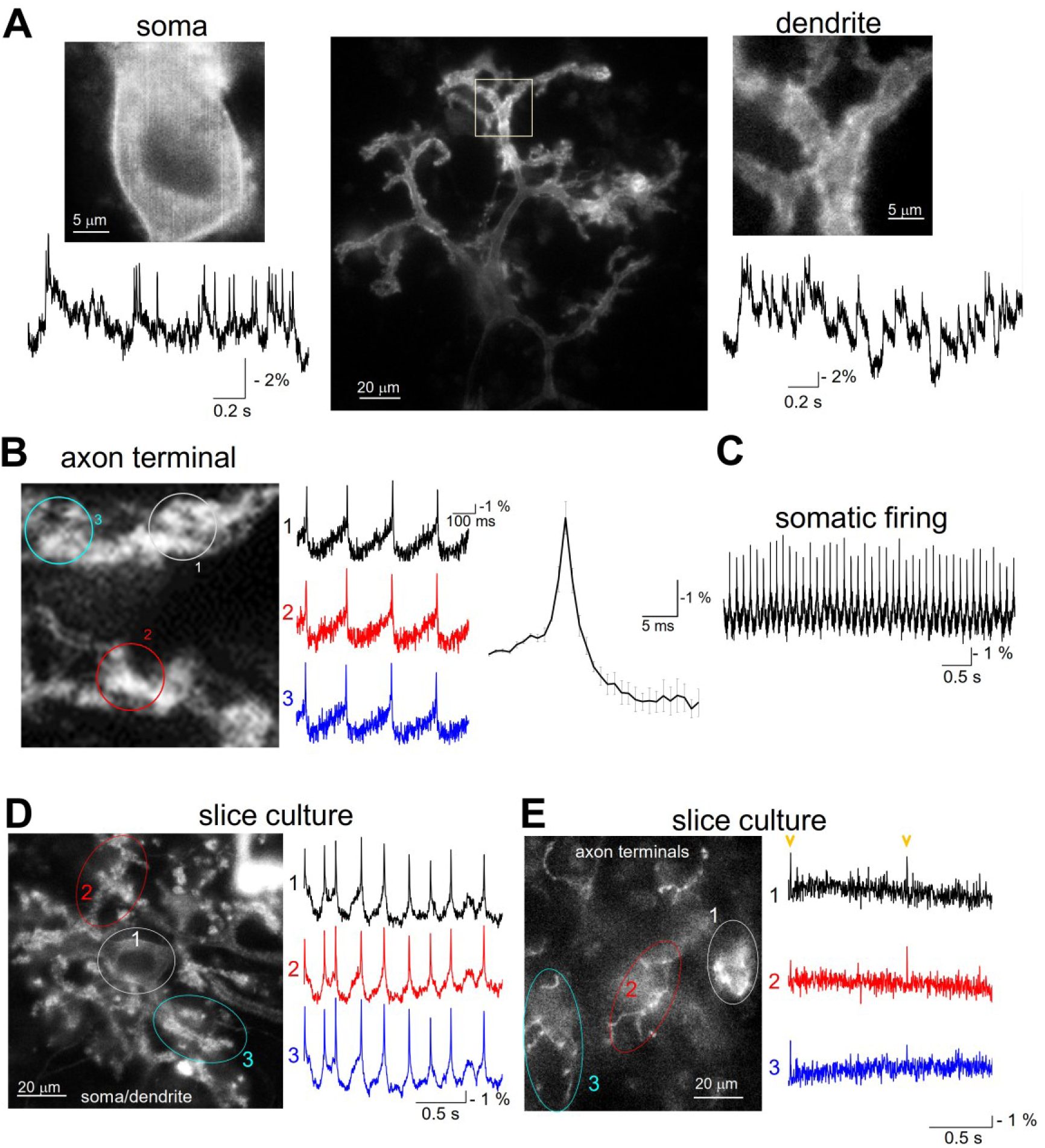
Imaging of APs and synaptic responses at PC sub-compartments. **A**, High-spatiotemporal axASAP3 voltage imagings of somatic AP firing and synaptic potentials recorded at the soma or dendrite of a cultured PC. **B**, Simultaneous multi-site voltage imaging along a long intact PC axon arbor. Traces 1–3 represent fluorescence changes recorded at different boutons along the same axon. Right trace is the average of fluorescently detected APs. **C**, Representative optical trace of spontaneous regular AP firings in a PC somatically recorded with axASAP3. **D**, **E**, Representative voltage imaging of APs and PSPs at PC soma/dendrite (D) and axon terminals (E) in an organotypic slice culture preparation, demonstrating robust detection of axonal spikes and subthreshold potentials in intact tissue arbors.

**Supplementary Fig. 4.**
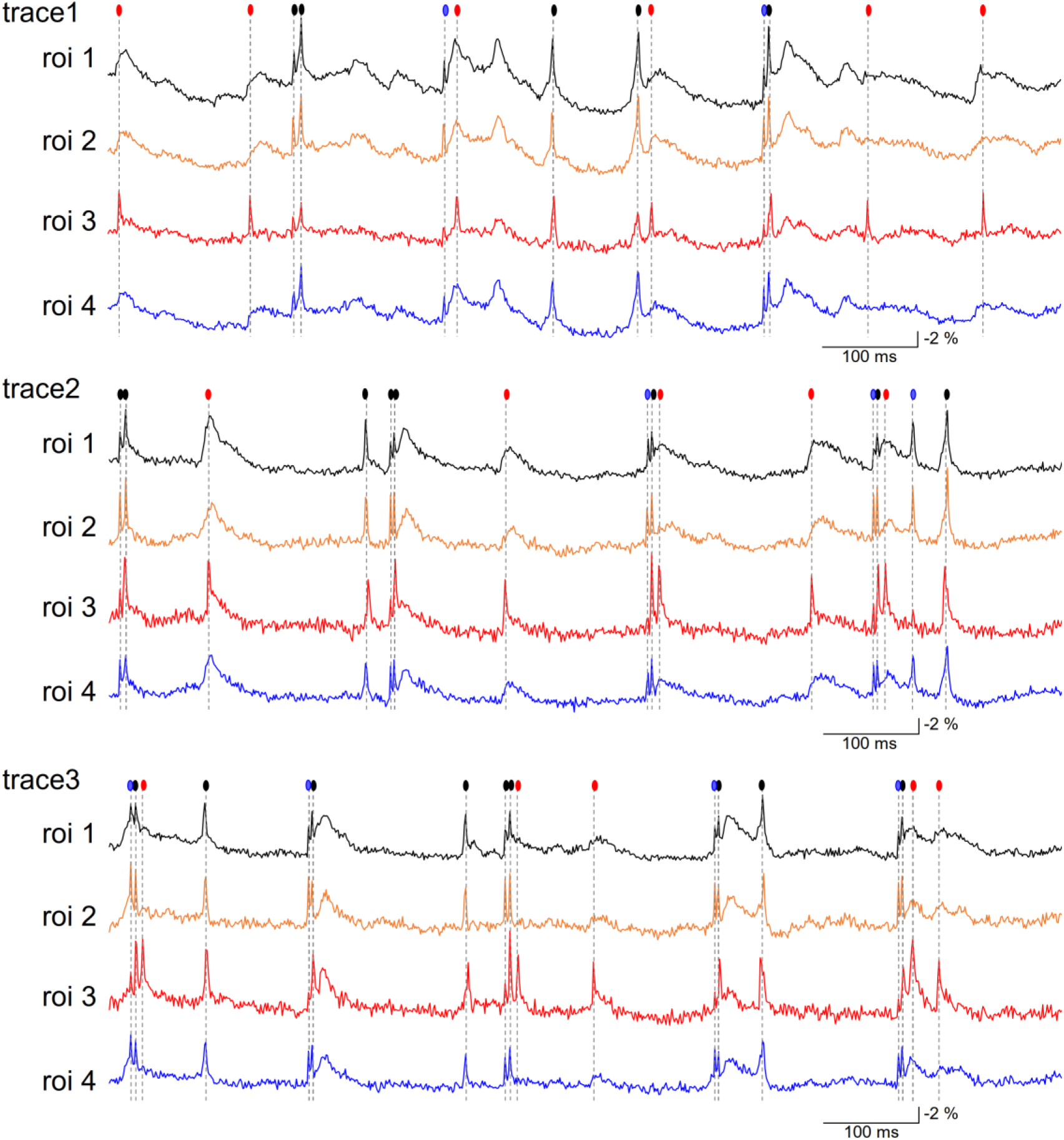
Regional heterogeneity and distinct firings along PC axonal arbors. Fluorescence time series recorded simultaneously from 4 distinct regions of interest (ROIs 1–4, indicated in Fig. 4D) along the axonal arbor of an identical PC 24 hours after the 2-min K-glu conditioning treatment. Three representative consecutive imaging trials (Trace 1-3) are shown. Filled circles above the traces indicate individual detected AP firing events: black circles denote global APs propagated faithfully to all 4 ROIs; blue, APs detected at 3 ROIs; orange, APs restricted to 2 ROIs; red, branch-specific ectopic AP firings detected uniquely at a single ROI (ROI 3). Note the occurrence of localized, non-propagating or branch-specific spikes following axonal plasticity.

**Supplementary Fig. 5.**
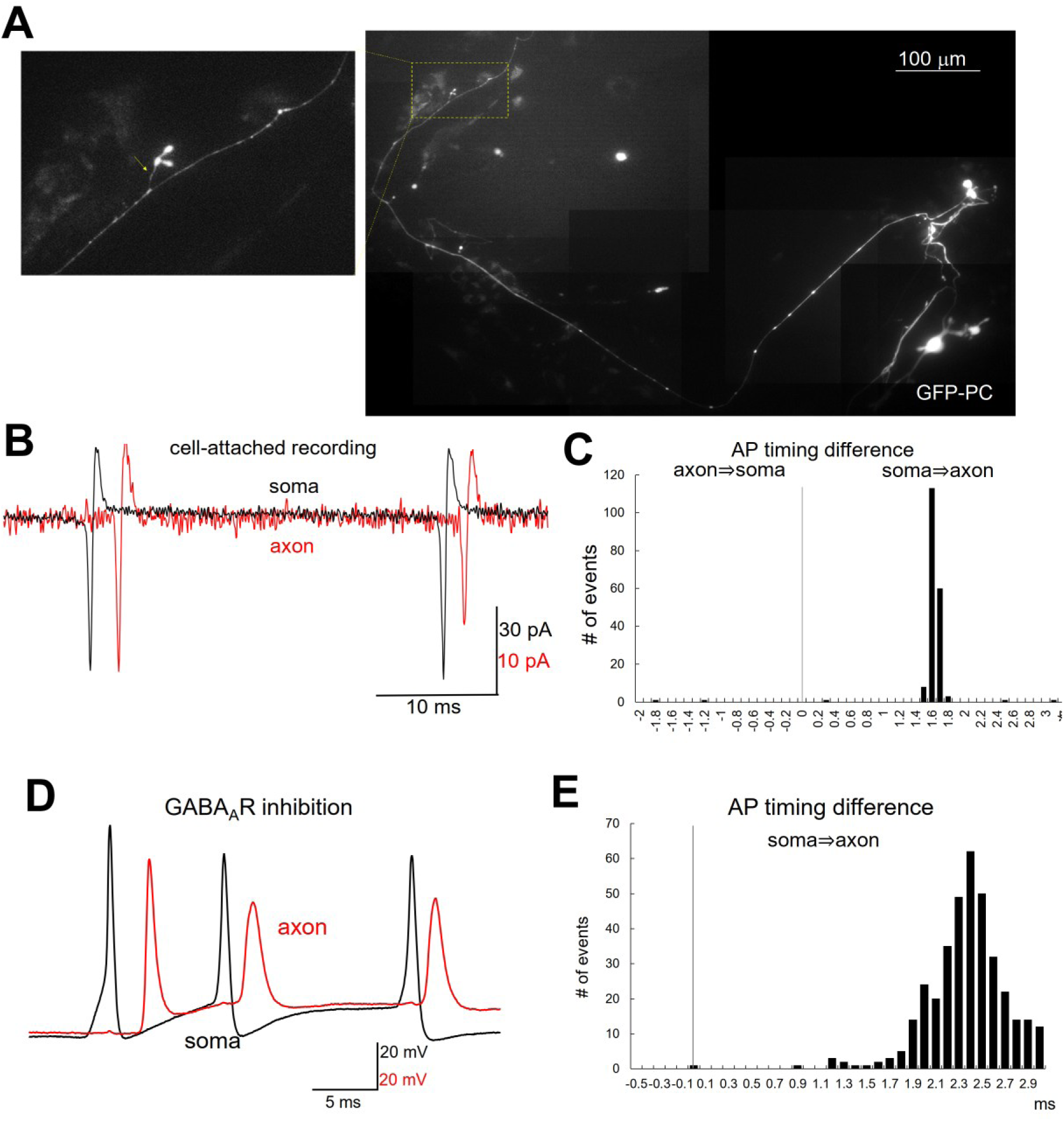
Spike initiation properties and propagation latency in control PCs. **A**, Representative fluorescence image of a GFP-labeled cultured PC subjected to dual cell-attached/whole-cell patch-clamp recordings from the soma and a distant axon terminal (enlarged images for the cell in Fig. 5A). Scale bar, 100 μm. **B**, Dual cell-attached current traces recorded simultaneously from the soma (black) and distal bouton (red) in a control PC, showing faithful somatic AP initiation prior to axonal propagation. **C**, Histogram of AP timing latency (Δt = t_bouton_ - t_soma_) between the soma and distal bouton in a representative control PC. Note the sharp, single-peak distribution centered around ∼2 ms, corresponding to forward orthodromic AP propagation from the soma/AIS to the distal terminal at a velocity of ∼0.5 m/s. **D**, **E**, Representative voltage traces (D) and latency histogram of spontaneous APs in control PCs after the application of 50 μM picrotoxin (GABA_A_R blocker) in a control PC. GABAergic inhibition did not affect the predominant somatic AP initiation in naive PCs.

**Supplementary Fig. 6.**
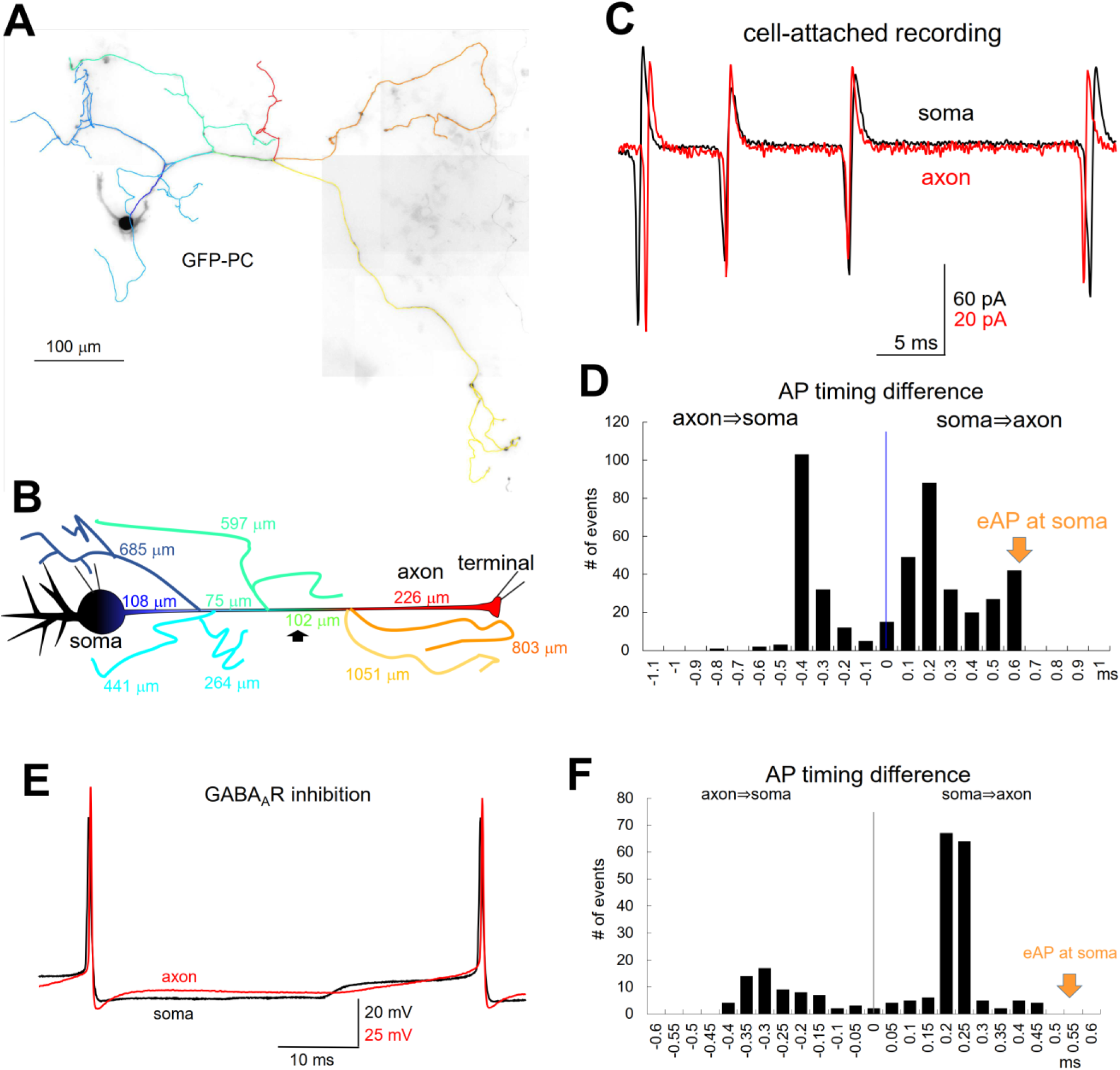
Anatomical matching between axonal branch lengths and multimodal firing latency distributions after K-glu-induced plasticity. **A**, Fluorescent image of a GFP-expressing PC 24 hours after the 2-min K-glu treatment, undergoing dual soma-bouton patch-clamp recording, with colored tracking of axonal arbors. **B**, Schematic morphological reconstruction with traced axonal arbor of the PC shown in A. Physical lengths (μm) between individual branch points in the main path from the soma to the recorded terminal, and those of total arbors bifurcated from the main path, are explicitly mapped (108 μm, 75 μm, 102 μm, 226 μm, 803μm, 1051μm, etc.). **C**, Representative dual cell-attached AP traces recorded simultaneously from the soma (black) and the designated distal terminal (red) 24 hours after the K-glu treatment. Note the occurrence of ectopic APs where axonal spikes precede somatic spikes. **D**, Histogram of AP timing difference (Δt) in the individual PC shown in A. Note the broad, multimodal peak distribution spanning both positive (soma to axon) and negative (axon to soma) values. Discrete peak intervals match the calculated conduction times to specific anatomical bifurcations mapped in B. **E**, **F**, Simultaneous current-clamp recordings (E) and latency distribution of AP initiation (F) from soma and terminal 24 hours after K-glu treatment in the presence of 50 μM picrotoxin. The persistence of ectopic negative-latency spikes indicates that increased intrinsic axonal membrane excitability primarily drives distal spike initiation, while local GABAergic axEPSPs further modulate local firing probability.

## Notes

### Competing Interest Statement

The authors have declared no competing interest.

